# Nonsynonymous mutations in *fepR* are associated with *Listeria* adaptation to low concentrations of benzalkonium chloride, but not increased survival in use level concentrations of benzalkonium chloride

**DOI:** 10.1101/2022.01.24.477644

**Authors:** Samantha Bolten, Anna Sophia Harrand, Jordan Skeens, Martin Wiedmann

**Affiliations:** Cornell University, Department of Food Science, Ithaca, NY

**Keywords:** *Listeria*, benzalkonium chloride, resistance, survival, adaptation, acclimation

## Abstract

Presence of and selection for *Listeria* spp. and *Listeria monocytogenes* that are less effectively inactivated by quaternary ammonium compounds (such as benzalkonium chloride [BC]) is a food safety concern, including in fresh produce environments. An initial MIC assay on 67 produce-associated *Listeria* strains showed that strains carrying BC resistance genes *bcrABC* (n=10) and *qacH* (n=1) showed higher MIC (4-6 mg/L BC) compared to strains lacking these genes (MIC of 1-2 mg/L BC). Serial passaging experiments that exposed the 67 strains to increasing BC concentrations revealed that 62/67 strains showed growth in BC concentrations above their parent MIC (range of 4-20 mg/L). Two serially passaged isolates were obtained for each parent strain and substreaked onto BHI agar in the absence of BC for seven rounds; 105/134 substreaked isolates showed higher substreaked MIC (range of 4 – 6 mg/L) compared to parent MIC. These results suggested isolates acquired genetic adaptations that confer BC resistance. Substreaked isolates were characterized by a combination of whole genome sequencing and Sanger sequencing of *fepR,* a local repressor of the MATE family efflux pump FepA. These data identified nonsynonymous *fepR* mutations in 48/67 isolates including 24 missense, 16 nonsense, and 8 frameshift mutations. The mean inactivation of substreaked isolates after exposure to use level concentrations of BC (300 mg/L) was 4.48 log, which was not significantly different from inactivation observed in parent strains. Serial passage experiments performed on cocultures of *Listeria* strains containing *bcrABC* or *qacH* did not yield growth at higher BC concentrations than monoculture experiments.

**IMPORTANCE:** *Listeria* resistance to quaternary ammonium compounds has been raised as a concern with regards to *Listeria* persistence in food environments, which can increase risk of product contamination, food recalls and foodborne illness outbreaks. Findings from our study show that individual *Listeria* strains can acquire genetic adaptations that confer resistance to low concentrations of benzalkonium chloride, but these genetic adaptations don’t increase *Listeria* survival when exposed to concentrations of benzalkonium chloride used for food contact surface sanitation (300 mg/L). Our study also suggests that there is limited risk of benzalkonium chloride resistance genes *bcrABC* and *qacH* spreading through horizontal gene transfer and conferring enhanced resistance of *L. monocytogenes* to benzalkonium chloride. Overall, our study suggests that emergence of benzalkonium chloride resistant *Listeria* strains in food processing environments is of limited concern, as even strains adapted to gain higher MIC *in vitro* maintain full sensitivity to use level concentrations of benzalkonium chloride.

## INTRODUCTION

*Listeria monocytogenes* is a foodborne pathogen that causes approximately 1600 cases of foodborne illness and 260 deaths per year in the US (1). *L. monocytogenes* persistence in built environments used for food production can provide a source of contamination in food products, which increases risk of recalls and foodborne illness outbreaks (2, 3). From 2009-2021, *L. monocytogenes* has been implicated in several outbreaks related to environmental contamination in fresh produce packing or processing environments, including outbreaks associated with celery (4), cantaloupe (5), caramel apples (6), and packaged lettuce (7).

Effective sanitation in the fresh produce supply chain is essential for mitigating *Listeria* contamination of fresh produce. Quaternary ammonium compounds are sanitizers that are used for sanitation of surfaces, equipment, and utensils in food production environments. Alkyl-(C_8_-C_18_) dimethyl benzyl ammonium chlorides (also known as benzalkonium chlorides) are a class of quaternary ammonium compounds that are approved for use on food contact surfaces at concentrations between 150-400 mg/L (termed henceforth as use level concentrations) in food processing and manufacturing environments (21 CFR 178.1010), and specific formulations of benzalkonium chlorides are widely used to control microorganisms such as *Listeria* in fresh produce packing and processing environments (8). There is concern that frequent use of benzalkonium chloride (BC) in food processing environments may be contributing to the selection of *Listeria* that are less effectively inactivated by BC in the food processing environment, which could increase the risk of *Listeria* persistence in food facilities (9, 10).

Genetically encoded efflux pump systems represent an important mechanism of BC resistance in *Listeria* (11). Two genes known to confer *Listeria* resistance to BC are *bcrABC* and *qacH* (12). Both *bcrABC* and *qacH* can be encoded on mobile genetic elements; the *bcrABC* cassette is most often found encoded on a pLM80-type plasmid (13, 14), and *qacH* is encoded on the *Tn6188* transposon (15). Additionally, both *bcrABC* and *qacH* have been frequently observed in *Listeria* strains isolated from food production environments (9, 12, 16–18). Because BC resistance genes are often encoded on mobile genetic elements, there is concern that horizontal gene transfer of these genes could facilitate widespread and enhanced resistance of *Listeria* to BC in the processing environment (19). Recently, horizontal gene transfer of a mobile genetic element containing *bcrABC* as well as cadmium resistance genes has been shown experimentally from *Listeria* species to a streptomycin resistant *L. monocytogenes*; in these experiments selection for streptomycin resistance and cadmium resistance was used to obtain transformants (20).

There is considerable debate surrounding the terminology used to describe the ability of *Listeria* to survive or grow in the presence of different concentrations of sanitizers like BC. With respect to clinical antibiotics, the term resistance is generally referred to as the ability of bacteria to maintain growth in therapeutic levels of an antibiotic agent (21). In the context of a food processing environment, improper development or implementation of sanitation standard operating procedures can allow for bacteria to be exposed to and potentially grow in the presence of sanitizers at concentrations below use level concentrations (10). Thus, in this study we sought to investigate the capacity of *Listeria* develop resistance to (or be able to grow in) low levels of BC. Here, we will use the term “resistance” as it is defined with respect to clinical antibiotics to refer to the capacity of *Listeria* strains to acquire the ability to grow in low levels of BC (defined here as <20 mg/L) that they were once sensitive to.

This same definition of resistance is not applicable when evaluating use level concentrations of sanitizers, because the efficacy of a sanitizer is measured by its ability to inactivate a certain number of microorganisms in a defined period (e.g., a 5-log reduction within 30 s for non-halide food contact sanitizers), not through its capacity to inhibit growth of microorganisms (22, 23). Therefore, in addition to assessing *Listeria’s* ability to grow in low levels of BC, we also sought to evaluate the survival of *Listeria* when exposed to use level concentrations of BC in this study.

The purpose of this study was to investigate (i) the sensitivity of monocultures of *Listeria* to low levels of BC (<20 mg/L), measured by assessing the minimum inhibitory concentration (MIC) of BC that suppresses *Listeria* growth and (ii) the effect that *Listeria* resistance to low concentrations of BC has on the survival of *Listeria* when exposed to use level concentrations of BC. Additionally, we also examined (iii) whether cocultures of *L. monocytogenes* carrying BC resistance genes *bcrABC* and *qacH* could acquire resistance to BC, through horizontal gene transfer, that exceeds resistance observed in monocultures.

## MATERIALS and METHODS

### Bacterial isolates and culture preparation

For this study, a diverse set of 67 *Listeria* strains from different fresh produce-associated sources was assembled. The selected strains represent pre-harvest (n=9), post-harvest (n=49), and retail (n=9) sources associated with fresh produce, including environmental samples (e.g., soil, water, environmental swabs) as well as actual produce samples (Table 1). The 67 strains were selected from a larger collection of 588 produce isolates (24), which had been characterized by whole genome sequencing and initial screening for inactivation when exposed to use level concentrations of three sanitizers, including BC; these data were used to select diverse BC tolerant and sensitive strains in this study. Strains were initially stratified into the 10% most BC tolerant and sensitive strains based on the preliminary data. For *L. monocytogenes*, we also selected nine of the most prevalent clonal complexes (CC) within the entire isolate collection, as well as the three most common CCs associated with lineage III (to assure representation of lineage III, which is less frequent than lineages I and II), yielding 12 common CCs; for these CCs we identified strains that represented the top 10% most BC tolerant and sensitive strains where available, but randomly selected strains within CCs that did not include strains that represented the 10% most BC tolerant and sensitive strains. In addition, the top three most tolerant isolates and top three most sensitive isolates from the entire culture collection were also selected if not already included in the strain set. Finally, any *L. monocytogenes* strains that represented unique genotypes with regard to presence/absence of *bcrABC* and *qacH* were also included (e.g., we included strain FSL R12-0189, which carried *qacH*, but was not classified in the top 10% most tolerant or sensitive strains). With this strategy, the final *L. monocytogenes* set included 28 strains that represented lineages I, II, and III (10, 15, and 3, respectively) and 16 clonal complexes (CC4, CC5, CC6, CC9, CC19, CC29, CC37, CC155, CC193, CC204, CC268, CC369, CC388, CC434, CC1789, ST1861); 12 of the strains represented the 10% most BC tolerant strains, 11 of the strains represented the 10% most BC sensitive strains, and five of the strains did not fall into these categories but were selected because they represented either lineage III (n=3) or represented a unique genotype with regards to presence of *bcrABC* or *qacH* (n=2). For species of *Listeria* other than *L. monocytogenes* (i.e., *L. innocua*, *L. marthii*, *L. welshimeri*, *L. seeligeri*, *L. ivanovii*), we picked the top 10% most BC tolerant and sensitive strains within each species; in addition we included one *L. welshimeri* isolate that carried *bcrABC* and a single *L. marthii* that contained the gene *lmo1409*, which represented a unique genotype for *L. marthii* strains from the original culture collection. The final strain set for non-*L. monocytogenes Listeria* species included 14 *L. seeligeri*, 12 *L. innocua*, eight *L. welshimeri*, three *L. marthii* and two *L. ivanovii* strains (Table 1). More information about the strains, including available metadata, serotypes, and associated publications can be found on Food Microbe Tracker (http://www.foodmicrobetracker.com/login/login.aspx) under a strain’s isolate ID (e.g., FSL H9-0078, see Table 1) (25).

**Table 1.**
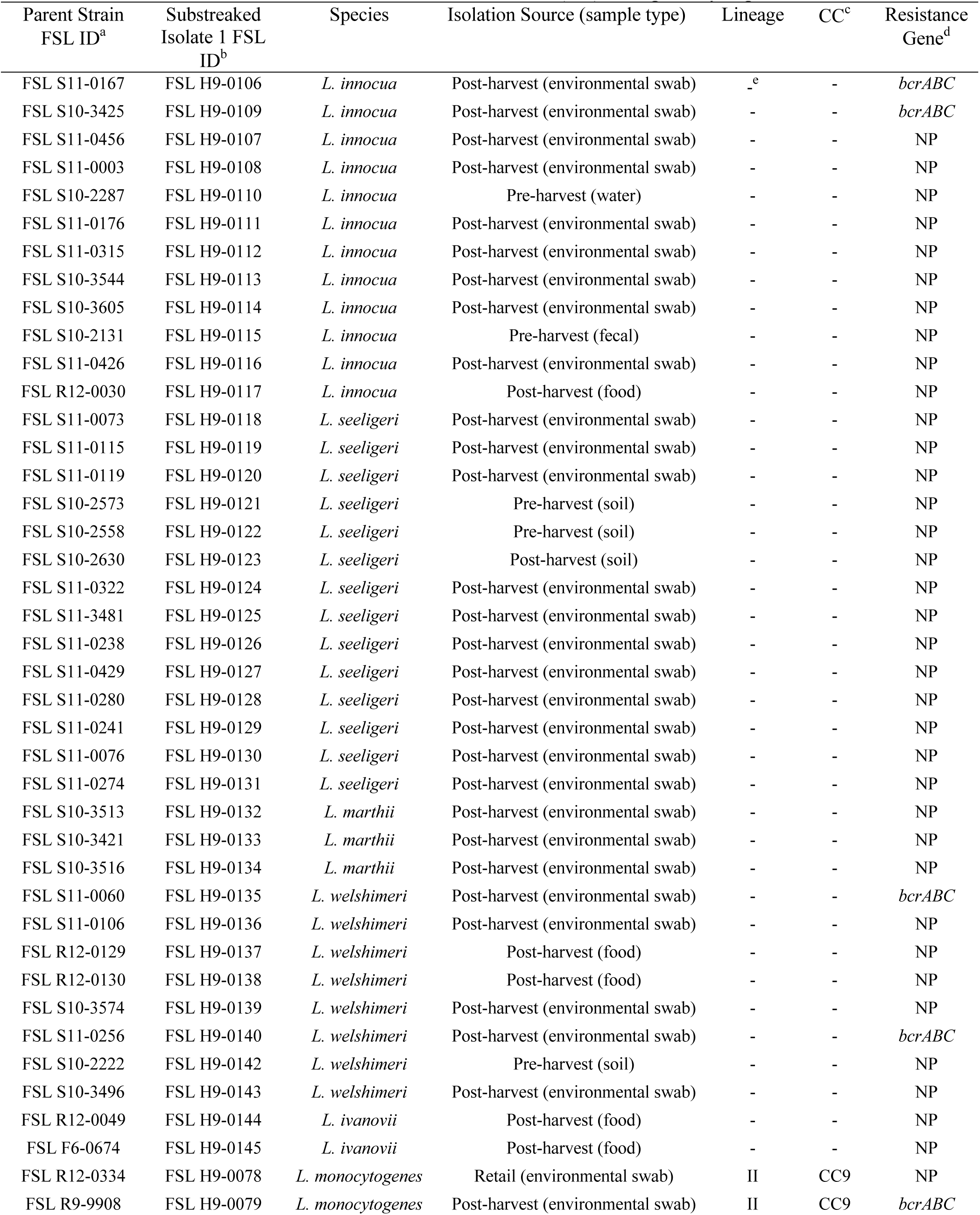
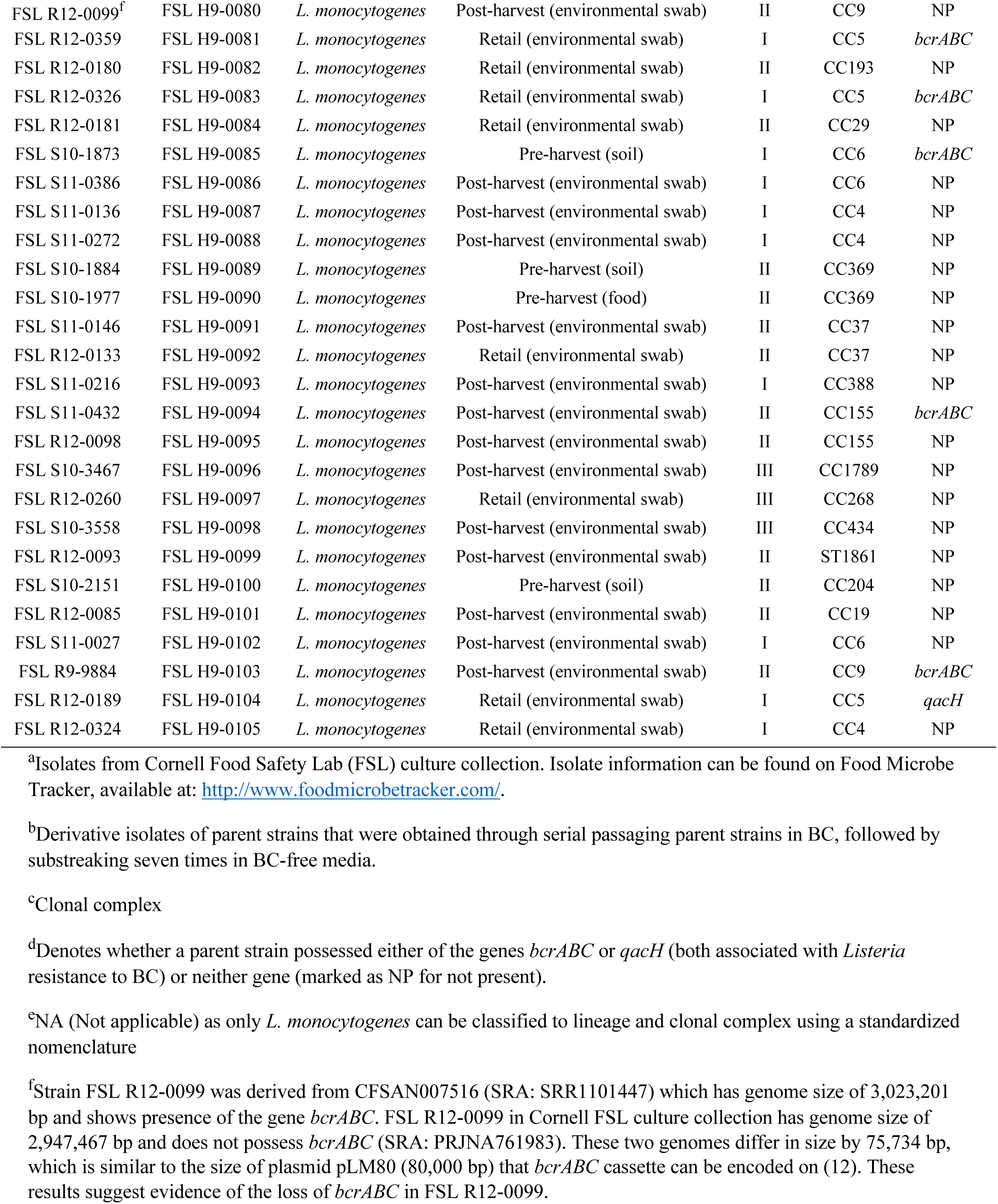
Listeria isolates selected for benzalkonium chloride (BC) susceptibility experiments.

| Parent Strain<br>FSL ID<sup>a</sup> | Substreaked<br>Isolate 1 FSL<br>ID<sup>b</sup> | Species | Isolation Source (sample type) | Lineage | CC<sup>c</sup> | Resistance<br>Gene<sup>d</sup> |
| --- | --- | --- | --- | --- | --- | --- |
| FSL S11-0167 | FSL H9-0106 | *L. innocua* | Post-harvest (environmental swab) | -<sup>e</sup> | - | *bcrABC* |
| FSL S10-3425 | FSL H9-0109 | *L. innocua* | Post-harvest (environmental swab) | - | - | *bcrABC* |
| FSL S11-0456 | FSL H9-0107 | *L. innocua* | Post-harvest (environmental swab) | - | - | NP |
| FSL S11-0003 | FSL H9-0108 | *L. innocua* | Post-harvest (environmental swab) | - | - | NP |
| FSL S10-2287 | FSL H9-0110 | *L. innocua* | Pre-harvest (water) | - | - | NP |
| FSL S11-0176 | FSL H9-0111 | *L. innocua* | Post-harvest (environmental swab) | - | - | NP |
| FSL S11-0315 | FSL H9-0112 | *L. innocua* | Post-harvest (environmental swab) | - | - | NP |
| FSL S10-3544 | FSL H9-0113 | *L. innocua* | Post-harvest (environmental swab) | - | - | NP |
| FSL S10-3605 | FSL H9-0114 | *L. innocua* | Post-harvest (environmental swab) | - | - | NP |
| FSL S10-2131 | FSL H9-0115 | *L. innocua* | Pre-harvest (fecal) | - | - | NP |
| FSL S11-0426 | FSL H9-0116 | *L. innocua* | Post-harvest (environmental swab) | - | - | NP |
| FSL R12-0030 | FSL H9-0117 | *L. innocua* | Post-harvest (food) | - | - | NP |
| FSL S11-0073 | FSL H9-0118 | *L. seeligeri* | Post-harvest (environmental swab) | - | - | NP |
| FSL S11-0115 | FSL H9-0119 | *L. seeligeri* | Post-harvest (environmental swab) | - | - | NP |
| FSL S11-0119 | FSL H9-0120 | *L. seeligeri* | Post-harvest (environmental swab) | - | - | NP |
| FSL S10-2573 | FSL H9-0121 | *L. seeligeri* | Pre-harvest (soil) | - | - | NP |
| FSL S10-2558 | FSL H9-0122 | *L. seeligeri* | Pre-harvest (soil) | - | - | NP |
| FSL S10-2630 | FSL H9-0123 | *L. seeligeri* | Post-harvest (soil) | - | - | NP |
| FSL S11-0322 | FSL H9-0124 | *L. seeligeri* | Post-harvest (environmental swab) | - | - | NP |
| FSL S11-3481 | FSL H9-0125 | *L. seeligeri* | Post-harvest (environmental swab) | - | - | NP |
| FSL S11-0238 | FSL H9-0126 | *L. seeligeri* | Post-harvest (environmental swab) | - | - | NP |
| FSL S11-0429 | FSL H9-0127 | *L. seeligeri* | Post-harvest (environmental swab) | - | - | NP |
| FSL S11-0280 | FSL H9-0128 | *L. seeligeri* | Post-harvest (environmental swab) | - | - | NP |
| FSL S11-0241 | FSL H9-0129 | *L. seeligeri* | Post-harvest (environmental swab) | - | - | NP |
| FSL S11-0076 | FSL H9-0130 | *L. seeligeri* | Post-harvest (environmental swab) | - | - | NP |
| FSL S11-0274 | FSL H9-0131 | *L. seeligeri* | Post-harvest (environmental swab) | - | - | NP |
| FSL S10-3513 | FSL H9-0132 | *L. marthii* | Post-harvest (environmental swab) | - | - | NP |
| FSL S10-3421 | FSL H9-0133 | *L. marthii* | Post-harvest (environmental swab) | - | - | NP |
| FSL S10-3516 | FSL H9-0134 | *L. marthii* | Post-harvest (environmental swab) | - | - | NP |
| FSL S11-0060 | FSL H9-0135 | *L. welshimeri* | Post-harvest (environmental swab) | - | - | *bcrABC* |
| FSL S11-0106 | FSL H9-0136 | *L. welshimeri* | Post-harvest (environmental swab) | - | - | NP |
| FSL R12-0129 | FSL H9-0137 | *L. welshimeri* | Post-harvest (food) | - | - | NP |
| FSL R12-0130 | FSL H9-0138 | *L. welshimeri* | Post-harvest (food) | - | - | NP |
| FSL S10-3574 | FSL H9-0139 | *L. welshimeri* | Post-harvest (environmental swab) | - | - | NP |
| FSL S11-0256 | FSL H9-0140 | *L. welshimeri* | Post-harvest (environmental swab) | - | - | *bcrABC* |
| FSL S10-2222 | FSL H9-0142 | *L. welshimeri* | Pre-harvest (soil) | - | - | NP |
| FSL S10-3496 | FSL H9-0143 | *L. welshimeri* | Post-harvest (environmental swab) | - | - | NP |
| FSL R12-0049 | FSL H9-0144 | *L. ivanovii* | Post-harvest (food) | - | - | NP |
| FSL F6-0674 | FSL H9-0145 | *L. ivanovii* | Post-harvest (food) | - | - | NP |
| FSL R12-0334 | FSL H9-0078 | *L. monocytogenes* | Retail (environmental swab) | II | CC9 | NP |
| FSL R9-9908 | FSL H9-0079 | *L. monocytogenes* | Post-harvest (environmental swab) | II | CC9 | *bcrABC* |
| FSL R12-0099<sup>f</sup> | FSL H9-0080 | *L. monocytogenes* | Post-harvest (environmental swab) | II | CC9 | NP |
| FSL R12-0359 | FSL H9-0081 | *L. monocytogenes* | Retail (environmental swab) | I | CC5 | *bcrABC* |
| FSL R12-0180 | FSL H9-0082 | *L. monocytogenes* | Retail (environmental swab) | II | CC193 | NP |
| FSL R12-0326 | FSL H9-0083 | *L. monocytogenes* | Retail (environmental swab) | I | CC5 | *bcrABC* |
| FSL R12-0181 | FSL H9-0084 | *L. monocytogenes* | Retail (environmental swab) | II | CC29 | NP |
| FSL S10-1873 | FSL H9-0085 | *L. monocytogenes* | Pre-harvest (soil) | I | CC6 | *bcrABC* |
| FSL S11-0386 | FSL H9-0086 | *L. monocytogenes* | Post-harvest (environmental swab) | I | CC6 | NP |
| FSL S11-0136 | FSL H9-0087 | *L. monocytogenes* | Post-harvest (environmental swab) | I | CC4 | NP |
| FSL S11-0272 | FSL H9-0088 | *L. monocytogenes* | Post-harvest (environmental swab) | I | CC4 | NP |
| FSL S10-1884 | FSL H9-0089 | *L. monocytogenes* | Pre-harvest (soil) | II | CC369 | NP |
| FSL S10-1977 | FSL H9-0090 | *L. monocytogenes* | Pre-harvest (food) | II | CC369 | NP |
| FSL S11-0146 | FSL H9-0091 | *L. monocytogenes* | Post-harvest (environmental swab) | II | CC37 | NP |
| FSL R12-0133 | FSL H9-0092 | *L. monocytogenes* | Retail (environmental swab) | II | CC37 | NP |
| FSL S11-0216 | FSL H9-0093 | *L. monocytogenes* | Post-harvest (environmental swab) | I | CC388 | NP |
| FSL S11-0432 | FSL H9-0094 | *L. monocytogenes* | Post-harvest (environmental swab) | II | CC155 | *bcrABC* |
| FSL R12-0098 | FSL H9-0095 | *L. monocytogenes* | Post-harvest (environmental swab) | II | CC155 | NP |
| FSL S10-3467 | FSL H9-0096 | *L. monocytogenes* | Post-harvest (environmental swab) | III | CC1789 | NP |
| FSL R12-0260 | FSL H9-0097 | *L. monocytogenes* | Retail (environmental swab) | III | CC268 | NP |
| FSL S10-3558 | FSL H9-0098 | *L. monocytogenes* | Post-harvest (environmental swab) | III | CC434 | NP |
| FSL R12-0093 | FSL H9-0099 | *L. monocytogenes* | Post-harvest (environmental swab) | II | ST1861 | NP |
| FSL S10-2151 | FSL H9-0100 | *L. monocytogenes* | Pre-harvest (soil) | II | CC204 | NP |
| FSL R12-0085 | FSL H9-0101 | *L. monocytogenes* | Post-harvest (environmental swab) | II | CC19 | NP |
| FSL S11-0027 | FSL H9-0102 | *L. monocytogenes* | Post-harvest (environmental swab) | I | CC6 | NP |
| FSL R9-9884 | FSL H9-0103 | *L. monocytogenes* | Post-harvest (environmental swab) | II | CC9 | *bcrABC* |
| FSL R12-0189 | FSL H9-0104 | *L. monocytogenes* | Retail (environmental swab) | I | CC5 | *qacH* |
| FSL R12-0324 | FSL H9-0105 | *L. monocytogenes* | Retail (environmental swab) | I | CC4 | NP |

<sup>a</sup>Isolates from Cornell Food Safety Lab (FSL) culture collection. Isolate information can be found on Food Microbe Tracker, available at: http://www.foodmicrobetracker.com/.
<sup>b</sup>Derivative isolates of parent strains that were obtained through serial passaging parent strains in BC, followed by substeaking seven times in BC-free media.
<sup>c</sup>Clonal complex
<sup>d</sup>Denotes whether a parent strain possessed either of the genes *bcrABC* or *qacH* (both associated with *Listeria* resistance to BC) or neither gene (marked as NP for not present).
<sup>e</sup>NA (Not applicable) as only *L. monocytogenes* can be classified to lineage and clonal complex using a standardized nomenclature
<sup>f</sup>Strain FSL R12-0099 was derived from CFSAN007516 (SRA: SRR1101447) which has genome size of 3,023,201 bp and shows presence of the gene *bcrABC*. FSL R12-0099 in Cornell FSL culture collection has genome size of 2,947,467 bp and does not possess *bcrABC* (SRA: PRJNA761983). These two genomes differ in size by 75,734 bp, which is similar to the size of plasmid pLM80 (80,000 bp) that *bcrABC* cassette can be encoded on (12). These results suggest evidence of the loss of *bcrABC* in FSL R12-0099.

Strains were streaked from -80°C freezer stocks (frozen in Brain Heart Infusion [BHI] broth with 15% glycerol) onto Brain Heart Infusion agar (BHIA; BD, Franklin Lakes, NJ), followed by incubation at 37°C for 24 h. Plates were stored at 4°C for at least 24 h and up to seven days for use in experiments. To prepare bacterial cultures, 5 mL of BHI broth was inoculated with a single colony, followed by static incubation at 22°C for 34 to 36 h; this culture was diluted 1:1,000 into fresh BHI broth, followed by static incubation at 22°C for 34 to 36 h to achieve bacterial cultures grown to stationary phase.

### MIC assay

The broth microdilution method was used to determine the MIC of individual strains to BC in a 96-well microtiter plate with 200 µL volume. BHI was supplemented with BC (n-alkyl dimethyl benzyl ammonium chloride, Sigma-Aldrich, St. Louis, MO, CAS: 63449-41-2) to achieve appropriate BC target concentrations. For the MIC assays, bacterial cultures grown to stationary phase as described above were diluted 1:1,000 and inoculated into BHI supplemented with BC (BHI-BC) to yield final BC concentrations of 0.25, 0.5, 1, 2, 4, 6, 8, 10, and 12 mg/L BC and a final bacterial concentration of ∼10^6^ CFU/mL per well. Absorbance of cultures was measured at 600 nm (OD_600_) in a microplate reader (Biotek Instruments, Winooski, VT) before (T0) and after incubation of 24 h at 22°C (T24). Differences in absorbance before and after 24 h incubation (ΔOD_600_) were calculated, and values >0.100 indicated strain growth at a given BC concentration.

### Serial passage experiment

Serial passaging was performed to assess *Listeria* capacity to acquire phenotypic resistance to BC. For serial passage experiments, each individual *Listeria* strain was serially passaged in increasing concentrations of BHI-BC in a 96-well microtiter plate with 200 µL volume. Bacterial cultures in stationary phase were diluted to a final concentration of ∼10^6^ CFU/mL at an initial concentration of 0.25 mg/L BHI-BC, followed by static incubation for up to 48 h at 22°C to allow for growth. Growth was assessed by measuring ΔOD_600_ at 24 h (as described above); strains were incubated for another 24 h if they did not show growth after 24 h. When growth was detected, strains were subcultured 1:10 in the next incremental concentration of BHI-BC (i.e., 0.5, 1, 2, with subsequent incremental increases of 2 mg/L BHI-BC). When strains failed to grow, the concentration a strain failed to grow in was diluted and dilutions were plated on BHIA, and incubated for 24 h at 37°C, for enumeration. The BC concentration at which strains failed to show growth was recorded as “serial passaged MIC”.

After serial passaging and plating onto BHIA, strains were assessed for the stability of their acquired resistance to BC. Two individual colonies from enumeration plates (defined as substreaked isolate 1 and substreaked isolate 2) from the serial passage experiment were substreaked onto BHIA in the absence of BC and incubated at 37°C for 24 h. This substreaking procedure was repeated for a total of seven times, and individual colonies of substreaked isolate 1 and substreaked isolate 2 from the seventh substreaked plate were assessed for their “substreaked MIC” as detailed above. The same colonies of substreaked isolate 1 and substreaked isolate 2 used for MIC experiments were frozen down in BHI with 15% glycerol and stored at -80°C.

### Coculture experiments

For coculture experiments, seven *L. monocytogenes* strains were conveniently selected to include strains with either *bcrABC*, *qacH*, or no resistance genes from lineages I and II (see Table 6). For each coculture experiment, four strains of *L. monocytogenes* were mixed together by adding 1 mL of prepared culture of each strain into a tube (referred to as cocultures), followed by thorough vortexing and centrifugation for 10 min at 4,000 rpm. Each coculture cell pellet was then resuspended in 200 µL BHI broth and the resuspension was transferred onto a 0.45 µM filter placed on a BHIA plate, followed by static incubation for 48 h at 22°C; this step was used to facilitate horizontal transfer of resistance genes. As a control, monocultures of each of the seven individual strains were plated in 200 µL volume on a 0.45 µM filter placed on BHIA and incubated for 48 h at 22°C. After incubation, each filter (including those for the monoculture controls) was removed with forceps and transferred into a 50 mL conical vial and washed with 5 mL BHI broth by pulse vortexing. Filters were aseptically removed, and filter rinsates were diluted to achieve OD_600_ of ∼0.2. OD-adjusted cultures were used to perform a MIC experiment (“filter plate MIC”) and were serially passaged in BC as described above. After serial passaging, three individual colonies from enumeration plates were substreaked seven times, and each of the three isolates was used to perform substreaked MIC experiments.

### Inactivation of parent strains and substreaked isolates of *Listeria* in use level concentration of BC

To determine whether substreaked isolates showed increased survival (compared to parent strains) in use level concentrations of BC, all 67 parent strains and all 67 substreaked isolate 1 isolates were exposed to use level concentrations of BC for 30 s. Prior to BC exposure, 1 mL of each bacterial culture was centrifuged at 10,000 g for 2 min to pellet cells. Pellets were resuspended in 1 mL of phosphate buffered saline adjusted to pH 8.0 (PBS-pH 8.0), and 200 µL aliquots of cell suspensions were subsequently transferred into individual wells of a 96-well deep well plate. Then, 200 µL of 600 mg/L BC (dissolved in PBS-pH 8.0) was added to each cell suspension, yielding a final exposure level of 300 mg/L BC; cells were exposed to this BC concentration for a total of 30 s, with pipetting up and down 10 times throughout the exposure period. Control cell suspensions were exposed to 200 µL PBS-pH 8.0 for 30 s. For neutralization, 400 µL of 1.43x Dey-Engley neutralizing broth (D/E broth; BD) was added by pipetting up and down 10 times, followed by incubation for 5 min. Neutralized cultures were serially diluted in D/E broth, followed by plating onto BHIA, incubation for 24 h at 37°C, and enumeration of colonies. Bacterial log reductions were calculated by subtracting the log CFU/mL of control cultures from log CFU/mL of BC treated cultures.

### Whole genome sequencing and bioinformatic analysis

Whole genome sequencing (WGS) was performed on 16 substreaked isolates; the resulting sequences were compared to sequences of parent strains to evaluate the presence of genetic mutations. The 16 isolates selected for WGS were randomly selected from the subset of 37 substreaked isolate 1 isolates that displayed a substreaked MIC at least 2-fold higher than their respective parent MIC. Single colonies of substreaked isolate 1 isolates were inoculated into 5 ml BHI broth, incubated for 15-18 h at 37°C, then DNA from cultures was extracted using the QiAamp DNA Mini Kit (Qiagen, Germantown, MD) following manufacturer’s instructions. Genomic DNA was sequenced using the Illumina NextSeq500 platform (Illumina, Inc. San Diego, CA) with 2×150 bp paired-end reads. The software Trimmomatic v 0.39 was used to trim adapters and low quality bases from raw sequencing data, and quality assessment was performed using FastQC v 0.11.8 (http://www.bioinformatics.babraham.ac.uk/projects/fastqc) (26). Assemblies were assembled using SPAdes v 3.15.2 using careful mode. Quality control of assemblies was performed using QUAST v 5.0.2 (27) and average coverage was determined using SAMtools v 1.11 (28). Genomes with an average coverage greater than 40x were included in genomic analysis. Contigs smaller than 500 bp were removed.

Genomes of substreaked isolates were compared with their respective parental genome through high quality SNP analysis (hqSNP) using the CFSAN SNP Pipeline v 2.2.1 (29). For each parent strain-substreaked isolate pair, the CFSAN SNP Pipeline was run twice; once aligning the paired-end reads of the substreaked isolate to the complete genome assembly of the parent strain, and once aligning the paired-end reads of the parent strain to the complete genome assembly of the substreaked isolate. SNPs that were identified in both analyses represented reliably identified SNP differences and were reported. If SNPs were located in an open reading frame the sequence of the open reading frame was searched in NCBI against the non-redundant protein sequences (nr) database using the BLASTX program to determine gene identities and/or their putative associated protein functions (30). SNPs were classified as synonymous (i.e., silent mutations) or nonsynonymous (i.e., missense, nonsense, or frameshift mutations) in Geneious Prime v 2021.1.1.

### PCR and Sanger sequencing

Mutations in *fepR* in substreaked isolates were identified through PCR amplification followed by Sanger sequencing of *fepR*. For lysate preparation, individual colonies of substreaked isolate 1 isolates were resuspended in 100 µL distilled H_2_O, and DNA was extracted by heat lysis via incubation at 95°C for 15 min. After incubation, the suspension was centrifuged at 14,000 g for 10 min, and supernatant was used as template for PCR. Two conventional PCR assays were developed and performed using *de novo* designed primers (Table 2) for amplification of 886 bp (for *L. monocytogenes*, *L. innocua*, and *L. marthii* isolates) and 819 bp (for *L. seeligeri* and *L. welshimeri* isolates) PCR products that contained the sequence of *fepR* (585 bp). PCR was conducted on an ABI 2720 thermal cycler (Thermo Fisher), using GoTaq Green Master Mix (Promega) with an initial denaturation of 5 min at 95°C, followed by 30 cycles of denaturation for 20 s at 95°C, annealing for 30 s at 55°C, and extension for 1 min at 72°C, and a final extension of 10 min at 72°C. Remaining primers and dNTPs were digested by adding exonuclease I (10 U, Thermo Fisher) and shrimp alkaline phosphatase (1 U, Thermo Fisher) to PCR products, and incubating samples at 37°C for 45 min, followed by 80°C for 15 min. Sanger sequencing was performed on PCR products by the Biotechnology Resource Center (Cornell University, Ithaca, NY). Alignments of *fepR* for each parent strain and its corresponding substreaked isolate 1 were performed in Geneious Prime v 2021.1.1.

**Table 2.**
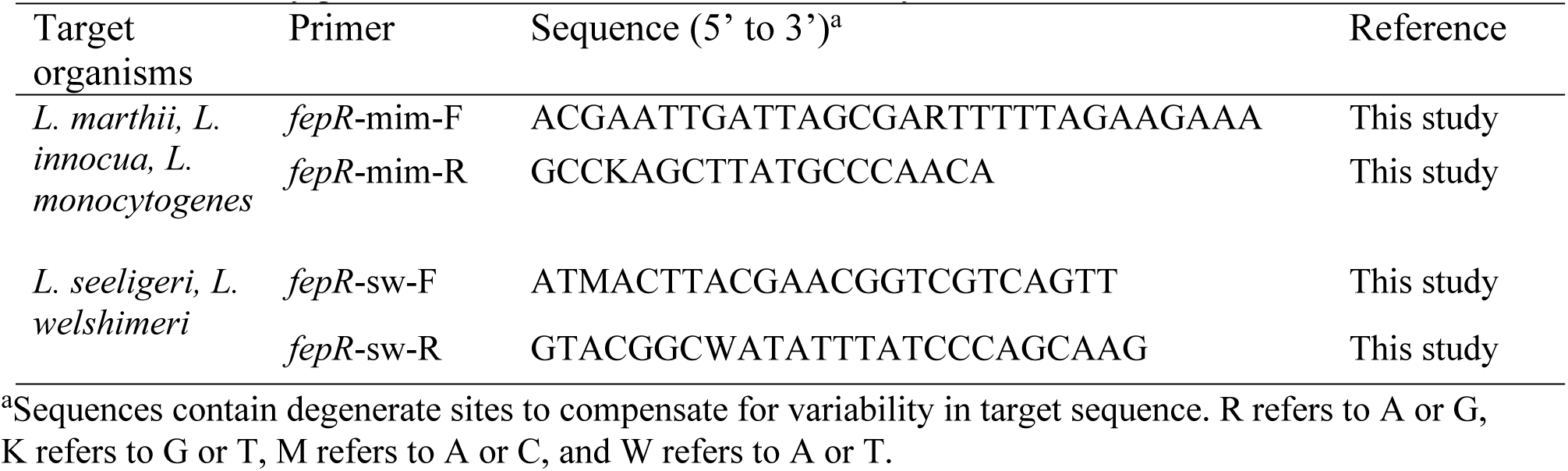
Primers for fepR in Listeria that were used in this study.

PCR amplification of *bcrABC* and *qacH* in parent strains and substreaked isolates was performed using primers and conditions described previously in (12).

### Statistical analysis

Data were analyzed in R, version 4.0.2 (R Core Team). MIC values were log transformed to satisfy normality assumptions. Linear mixed effects models were fit using the lme4 package to determine (i) the effect of the interaction between type of MIC (i.e., parent, serial passaged, substreaked) and whether an isolate was *L. monocytogenes* or *L.* spp. (i.e., *L. innocua*, *L. seeligeri*, *L. welshimeri*, *L. marthii*, or *L. ivanovii*), on the outcome of MIC value, (ii) the effect of the interaction between type of MIC, and whether an isolate carried a BC resistance gene (*bcrABC* or *qacH*), on the outcome of MIC value, (iii) the effect of whether a substreaked isolate showed a mutation in *fepR* on the outcome of MIC value, and (iv) the effect of type of MIC, and whether *L. monocytogenes* strains were grown in cocultures or monocultures, on the outcome of MIC value. For these analyses, type of MIC, *L. monocytogenes* vs *L.* spp., and resistance gene (presence or absence) were considered fixed effects, and each individual isolate’s FSL ID was considered a random effect. Analysis of variance (ANOVA) was performed on linear mixed-models, followed by post-hoc analysis of estimated marginal means with Tukey adjustment using the emmeans package in R (31).

To assess the effect of exposure to use level concentrations of BC on the log reductions of parent strains compared to substreaked isolates, an unpaired t-test was used to compare the mean log reductions of all parent strains to the mean log reductions of all substreaked isolates. One-way ANOVA was also used to assess for the association of BC resistance gene presence on the outcome of log reduction following exposure to use level concentrations of BC. P values of ≤0.05 were considered statistically significant. The detection limit for bacterial colony enumeration was 2 log CFU/mL. In cases where no colonies were observed, the value for the limit of detection (2 log CFU/mL) was used for statistical analyses. Raw data and the R code are available on GitHub (https://github.com/sjb375/Listeria_BC_adaptation).

## RESULTS

### Parent strains showed MIC values to BC ranging from 1 to 6 mg/L, with strains carrying BC resistance genes showing significantly higher MIC values

Initial MIC experiments on the 67 *Listeria* strains characterized here showed a narrow range of MIC values ranging from 1-6 mg/L with an estimated marginal mean of 2.30 mg/L (Figure 1); these initial MIC values will be referred to as “parent MIC”. Results from two-way ANOVA and post-hoc tests (Table 3) showed that *Listeria* strains that carry the BC resistance genes *bcrABC* (n=10) or *qacH* (n=1) showed significantly higher MIC (estimated marginal mean of 5.37 mg/L; range of 4-6 mg/L BC) as compared to strains lacking these genes (estimated marginal mean MIC of 1.95 mg/L; range of 1-2 mg/L BC) (P<0.05). *bcrABC* was present in six *L. monocytogenes*, two *L. innocua*, and two *L. welshimeri* isolates; *qacH* was present in one *L. monocytogenes* strain. Parent MICs for *L. monocytogenes* and *Listeria* spp. (i.e., all *L. innocua*, *L. marthii*, *L. welshimeri*, *L. seeligeri*, and *L. ivanovii* strains examined in this study) were not significantly different from one another with estimated marginal mean MIC for *L. monocytogenes* of 2.59 mg/L (range of 2-6 mg/L) and estimated marginal mean MIC for *Listeria* spp. of 2.12 mg/L (range of 1-6 mg/L) (P>0.05).

**Figure 1.**
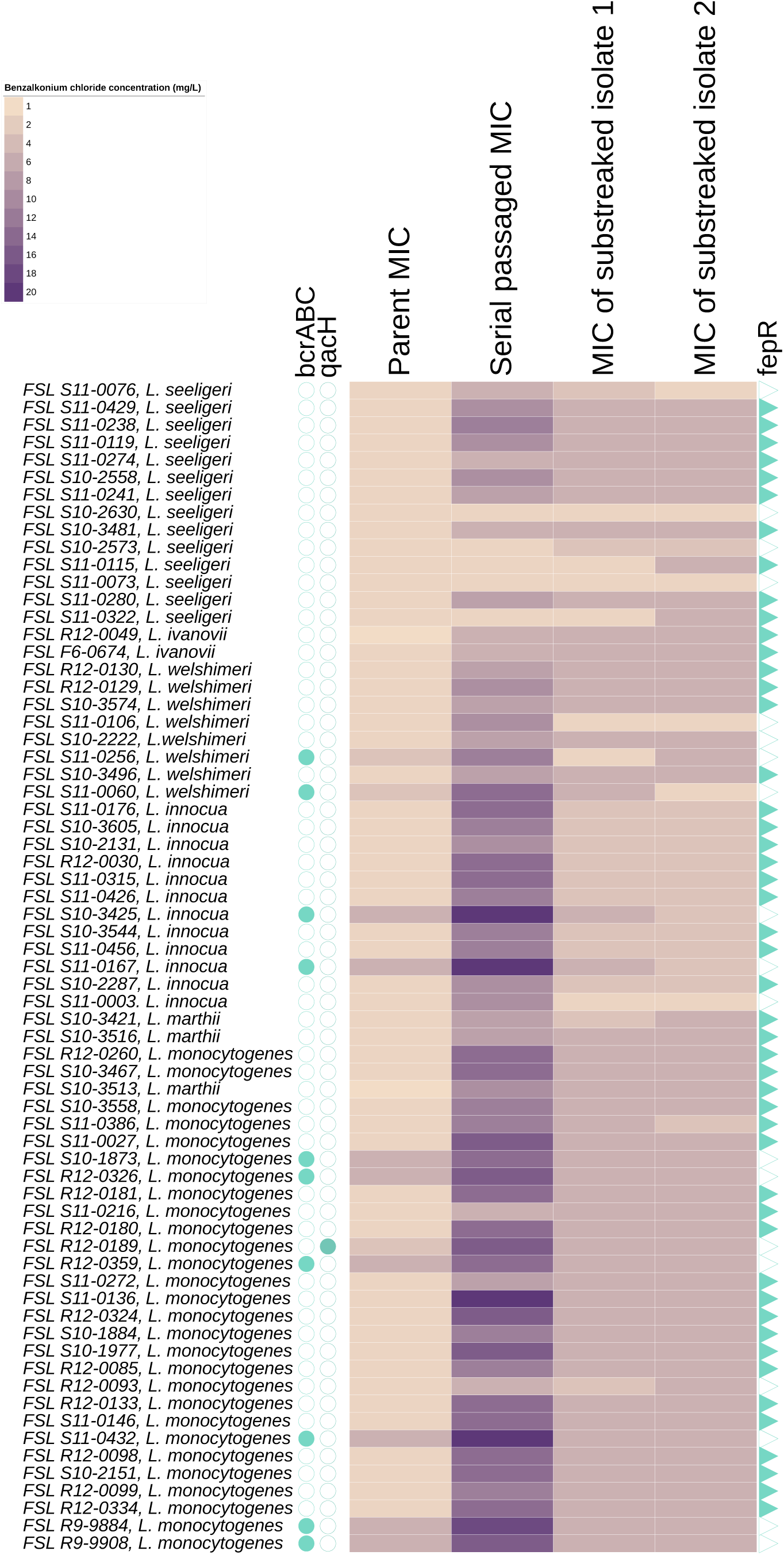
Heatmap of *Listeria* MIC to benzalkonium chloride (BC). Each row is specific to an individual parent strain of *Listeria*. For example, row 1 of the heatmap corresponds to strain FSL S11-0076, which is a strain of *L. seeligeri.* Parent MIC indicates the initial MIC obtained for a given *Listeria* strain; serial passaged MIC indicates the MIC obtained through serial passaging a strain in increasing concentrations of BC; MIC of substreaked isolate 1 and MIC of substreaked isolate 2 indicates the MIC obtained from two isolates obtained from serial passage experiments that were substreaked seven times onto BHI agar. Blue filled in circles represent the presence of a BC resistance gene (*bcrABC*, *qacH*) in the *Listeria* parent strain; FSL S11-0256 and FSL S11-0060 showed loss of *bcrABC* in their corresponding substreaked isolate 1, and FSL S11-0256 and FSL R9-9884 showed loss of *bcrABC* in their corresponding substreaked isolate 2. Blue filled in triangles represent the presence of a nonsynonymous mutation in *fepR* in substreaked isolate 1.

**Table 3.** Marginal means estimates for MIC values based on interaction models I and II.

| Interaction model | Fixed effects interaction | Levels | Estimate (mg/L) | Standard Error | Lower CL<sup>a</sup> | Upper CL | Group<sup>b</sup> |
| --- | --- | --- | --- | --- | --- | --- | --- |
| I<sup>c</sup> | MIC type: *Listeria* type | parent MIC : Lm | 2.59 | 0.19 | 2.22 | 2.97 | a |
|  |  | parent MIC : *L. spp* | 2.12 | 0.13 | 1.86 | 2.37 | a |
|  |  | serial passaged MIC : Lm | 13.38 | 0.98 | 11.45 | 15.31 | e |
|  |  | serial passaged MIC : *L. spp.* | 8.03 | 0.50 | 7.05 | 9.01 | d |
|  |  | substreaked MIC : Lm | 5.91 | 0.34 | 5.25 | 6.58 | c |
|  |  | substreaked MIC : *L. spp.* | 4.41 | 0.21 | 3.99 | 4.83 | b |
| II<sup>d</sup> | MIC type: Resistance gene | parent MIC : present | 5.37 | 0.59 | 4.21 | 6.53 | b |
|  |  | parent MIC : absent | 1.95 | 0.10 | 1.76 | 2.14 | a |
|  |  | serial passaged MIC : present | 16.14 | 1.77 | 12.66 | 19.62 | d |
|  |  | serial passaged MIC : absent | 9.04 | 0.44 | 8.17 | 9.90 | c |
|  |  | substreaked MIC : present | 5.23 | 0.46 | 4.32 | 6.14 | b |
|  |  | substreaked MIC : absent | 4.94 | 0.19 | 4.56 | 5.32 | b |

<sup>a</sup>Confidence limit.
<sup>b</sup>Group refers to significant differences based on post-hoc multiple-comparison adjustment with Tukey's honestly significant differences (HSD) test. Within each interaction model (I or II) groups that do not share a given letter are significantly different (P<0.05).
<sup>c</sup>Interaction model I tests the interaction of fixed effects of (i) *Listeria* type (either Lm (*L. monocytogenes*) or *L. spp* (*L. innocua*, *L. welshimeri*, *L. seeligeri*, *L. ivanovii*, and *L. marthii*)) and (ii) MIC type (either parent MIC, serial passaged MIC, or substreaked MIC) on MIC to benzalkonium chloride. Parent MIC: the initial MIC for a *Listeria* strain; serial passaged MIC: the MIC obtained through serial passaging a strain in increasing concentrations of benzalkonium chloride; substreaked MIC: the MIC obtained from two isolates obtained from serial passage experiments that were substreaked seven times onto BHI agar.
<sup>d</sup>Interaction model II tests the interaction of fixed effects of (i) Resistance gene present (*Listeria* strains carry *bcrABC* or *qacH*) or absent (*Listeria* strains that do not carry *bcrABC* or *qacH*) and MIC type (either parent MIC, serial passaged MIC, or substreaked MIC) on MIC values in benzalkonium chloride.

### Serial passaging experiments showed that 62/67 parent strains were able to grow in BC concentrations above their initial MIC

To assess the ability of *Listeria* to acquire increased resistance to BC, monocultures of all 67 parent strains of *Listeria* were subjected to serial passaging in increasing concentrations of BC. The highest concentration of BC where a given strain failed to show growth after 48 h incubation was designated as “serial passaged MIC”. The estimated marginal mean serial passaged MIC for all 67 strains was 9.94 mg/L, which was significantly higher than the estimated marginal mean parent MIC of 2.30 mg/L (P<0.05). Overall, 62/67 strains were able to grow to BC concentrations above their parent MIC; for these strains’ serial passaged MIC values ranged from 4-20 mg/L. The five strains that did not grow above parent MIC all belonged to the species *L. seeligeri*; *L. seeligeri* was represented by a total of 14 strains characterized in this study. Results from two-way ANOVA and post-hoc tests showed that the serial passaged MIC for *L. monocytogenes* (estimated marginal mean of 13.38 mg/L; range of 6-20 mg/L) was significantly higher (P<0.05) compared to the serial passaged MIC for *Listeria* spp. (estimated marginal mean of 8.03 mg/L; range of 2-20 mg/L) (Table 3).

To further characterize the impact of serial passages in the presence of BC on MICs, we also calculated the MIC fold change for all strains (Table 4). The fold change of serial passaged MIC/parent MIC for all strains ranged from 1 to 10. *L. monocytogenes* showed slightly higher fold change (estimated marginal mean fold change of 4.74) compared to *Listeria* spp. (estimated marginal mean fold change of 4.03), but this difference was not significant (P>0.05). The fold change of serial passaged MIC/parent MIC for *Listeria* strains that carried BC resistance genes *bcrABC* or *qacH* was significantly lower (estimated marginal mean fold change of 2.25) than the fold change observed in strains that did not carry these BC resistance genes (estimated marginal mean fold change of 4.90) (P<0.05).

**Table 4.** Marginal means estimates for fold change of MIC values based on additive models I and II.

| Model | Fixed effects | Levels | Estimated fold change | Standard error | Lower CL<sup>a</sup> | Upper CL | Group<sup>b</sup> |
| --- | --- | --- | --- | --- | --- | --- | --- |
| I<sup>c</sup> | Fold change type<br>*Listeria* type | Serial passaged MIC/parent MIC : Lm | 4.74 | 0.43 | 3.89 | 5.60 | b |
|  |  | Serial passaged MIC/parent MIC : *L. spp* | 4.03 | 0.32 | 3.40 | 4.66 | b |
|  |  | substreaked MIC/parent MIC : Lm | 2.38 | 0.21 | 1.97 | 2.79 | a |
|  |  | substreaked MIC/parent MIC : *L. spp* | 2.02 | 0.15 | 1.73 | 2.32 | a |
| II<sup>d</sup> | Fold change type<br>Resistance gene | Serial passaged MIC/parent MIC : present | 2.25 | 0.25 | 1.76 | 2.74 | b |
|  |  | Serial passaged MIC/parent MIC : absent | 4.90 | 0.28 | 4.35 | 5.46 | c |
|  |  | substreaked MIC/parent MIC : present | 1.13 | 0.12 | 0.89 | 1.36 | a |
|  |  | substreaked MIC/parent MIC : absent | 2.46 | 0.12 | 2.22 | 2.70 | b |

<sup>a</sup>Confidence limit.
<sup>b</sup>Group refers to significant differences based on post-hoc multiple-comparison adjustment with Tukey's honestly significant differences (HSD) test. Within each additive model (I or II) different numbers denote significant differences (P<0.05).
<sup>c</sup>Additive model I tests the fixed effects of *Listeria* type (either Lm (*L. monocytogenes*) or *L. spp* (*L. innocua*, *L. welshimeri*, *L. seeligeri*, *L. ivanovii*, and *L. marthii*)) and MICs being compared (serial passaged MIC/parent MIC, substreaked MIC/parent MIC) on the level of fold change. Parent MIC: the initial MIC for a *Listeria* strain; serial passaged MIC: the MIC obtained through serial passaging a strain in increasing concentrations of benzalkonium chloride; substreaked MIC: the MIC obtained from two isolates obtained from serial passage experiments that were substreaked seven times onto BHI agar.
<sup>d</sup>Additive model II tests the fixed effects of (i) Resistance gene present (*Listeria* strains carry *bcrABC* or *qacH*) or absent (*Listeria* strains that do not carry *bcrABC* or *qacH*) and MICs being compared (serial passaged MIC/parent MIC, substreaked MIC/parent MIC) on the level of fold change.

### *Listeria* maintain increased MIC values after substreaking seven rounds in the absence of BC

For each serial passage, the culture that represented the concentration of BHI-BC where a given *Listeria* failed to show growth was plated onto BC-free BHIA and two individual colonies (substreaked isolate 1 and substreaked isolate 2) were selected for further characterization. All colonies were substreaked for seven rounds on BC-free BHIA to determine whether *Listeria* resistance to BC was maintained in absence of selective pressure. These data were used to determine whether increased MIC values acquired during serial passaging were due to transient or inherited resistance. After seven rounds of substreaking, all isolates were characterized through MIC experiments; the resulting MIC values were designated as “MIC of substreaked isolate 1” and “MIC of substreaked isolate 2” (Figure 1). The estimated marginal mean substreaked MIC was 4.99 mg/L (range of 2-6 mg/L), which was significantly higher than the estimated marginal mean parent MIC (2.30 mg/L), but also significantly lower than the estimated marginal mean serial passaged MIC (9.94 mg/L) (P<0.05).

The majority of substreaked isolates showed higher substreaked MIC compared to their parent MIC (105/134), but lower substreaked MIC values compared to their serial passaged MIC (113/134). Results from two-way ANOVA and post-hoc tests showed that *L. monocytogenes* substreaked MIC values were significantly higher (estimated marginal mean of 5.91 mg/L, range of 4-6 mg/L) compared to *Listeria* spp. substreaked MIC values (estimated marginal mean of 4.41 mg/L, range of 2-6 mg/L) (P<0.05). The presence (estimated marginal mean of 5.23 mg/L, range of 2-6 mg/L) or absence (estimated marginal mean of 4.94 mg/L, range of 2-6 mg/L) of BC resistance genes *bcrABC* or *qacH* was not associated with a significant difference in substreaked MIC (P>0.05) (Table 3).

Further analysis of the fold change of substreaked MIC/parent MIC (Table 4) showed that the fold change of *L. monocytogenes* (estimated marginal mean fold change of 2.38) did not differ significantly from the fold change of *Listeria* spp. (estimated marginal mean fold change of 2.02) (P>0.05), suggesting no difference in the ability of *L. monocytogenes* and *Listeria* spp. to adapt to BC. The estimated marginal mean fold change of substreaked MIC/parent MIC for *Listeria* that carried either *bcrABC* or *qacH* was significantly lower (estimated marginal mean fold change of 1.13) compared to the fold change of isolates that did not carry *bcrABC* or *qacH* (estimated marginal mean fold change of 2.46) (P<0.05), suggesting that isolates carrying *bcrABC* or *qacH* showed limited adaptation to BC.

The 67 strains in this study were selected from a larger culture collection where they were previously classified as the top 10% tolerant, top 10% sensitive, or average in their sensitivity (representing the 11-89th percentile) to use level concentrations of BC (based on log reduction data collected in a previous study) (24). Hence, we examined whether these previous classifications for the *Listeria* strains we selected were associated with the MIC values for these strains. Our results found that the previous classification of *Listeria* strains as tolerant, average, or sensitive to use levels of BC was not significantly associated with a difference in parent MIC, serial passaged MIC, or substreaked MIC values (P>0.05).

### Mutations in transcriptional regulator gene *fepR* are associated with *Listeria* resistance to low levels of BC

To investigate putative mutations responsible for the observed phenotypic resistance in substreaked isolates, we performed WGS on a subset of substreaked isolate 1 isolates (*n*=16) and compared their genomes to the genomes of their respective parent strains using high quality SNP (hqSNP) analysis. In all 16 substreaked isolates sequenced, we detected mutations in *fepR* (lmo2088), a gene encoding a TetR family transcriptional regulator (Table 5). Mutations detected through hqSNP analysis included (i) nonsense mutations (i.e., SNPs that lead to premature stop codons in the coding sequence of *fepR*) in 8/16 isolates, and (ii) missense mutations (i.e., SNPs that lead to an amino acid change in the *fepR* coding sequence) in 5/16 isolates. In addition, single nucleotide deletions leading to frameshift mutations in *fepR* were detected in 3/16 isolates; these deletions were detected by aligning genome assemblies of parent strains and substreaked isolates to an annotated *fepR* sequence. Other mutations detected in substreaked isolates included a missense mutation in an internalin-like protein in substreaked isolate FSL H9-0100, and a nonsense mutation detected in the putative membrane protein YdfK in substreaked isolate FSL H9-0112.

**Table 5.** Mutations detected in *fepR* for substreaked isolates of *Listeria*. Mutations were identified by comparison of the whole genome sequences of parent strains and substreaked isolates.

| Substreaked Isolate ID | Species | Total number of SNPs in substreaked isolate | Mutation detected in *fepR*<sup>a</sup> | Mutation type<sup>b</sup> in *fepR* | BioProject Accession number or SRA ID of parent strain<sup>c</sup> |
| --- | --- | --- | --- | --- | --- |
| FSL H9-0080 | *L. monocytogenes* | 0 | Deletion | Frameshift | PRJNA761983 |
| FSL H9-0082 | *L. monocytogenes* | 1 | SNP | Missense | SRR9019233 |
| FSL H9-0095 | *L. monocytogenes* | 2 | SNP | Nonsense | SRR1027706 |
| FSL H9-0098 | *L. monocytogenes* | 2 | SNP | Missense | PRJNA761983 |
| FSL H9-0100 | *L. monocytogenes* | 4 | SNP | Nonsense | SRR12696480 |
| FSL H9-0097 | *L. monocytogenes* | 1 | SNP | Nonsense | SRR9019214 |
| FSL H9-0105 | *L. monocytogenes* | 1 | Deletion | Frameshift | SRR9019309 |
| FSL H9-0111 | *L. innocua* | 2 | SNP | Nonsense | PRJNA761983 |
| FSL H9-0112 | *L. innocua* | 4 | SNP | Nonsense | PRJNA761983 |
| FSL H9-0122 | *L. seeligeri* | 1 | SNP | Missense | PRJNA761983 |
| FSL H9-0131 | *L. seeligeri* | 2 | SNP | Nonsense | PRJNA761983 |
| FSL H9-0132 | *L. marthii* | 2 | SNP | Missense | PRJNA761983 |
| FSL H9-0134 | *L. marthii* | 1 | SNP | Missense | PRJNA761983 |
| FSL H9-0137 | *L. welshimeri* | 0 | Deletion | Frameshift | PRJNA761983 |
| FSL H9-0144 | *L. ivanovii* | 1 | SNP | Nonsense | PRJNA761983 |
| FSL H9-0145 | *L. ivanovii* | 2 | SNP | Nonsense | PRJNA761983 |

<sup>a</sup>*fepR* encodes for a transcriptional regulator in the TetR family of transcriptional regulators (GenBank: WP\_010991061.1). SNP refers to single nucleotide polymorphism detected in *fepR*, Deletion refers to a deletion of a single nucleotide *fepR*.
<sup>b</sup>Either missense (SNP confers amino acid shift in FepR protein), nonsense (either SNP confers a premature stop codon in FepR), or deletion (confers a frameshift in FepR).
<sup>c</sup>Whole genome sequences of all substreaked isolates can be found in BioProject Accession number: PRJNA761983. Parent strain ID that corresponds to substreaked isolate ID is shown in Table 1.

On the basis of the high frequency of *fepR* mutations in substreaked isolates, we PCR amplified the *fepR* sequence, followed by Sanger sequencing of PCR amplicons, for the 51 remaining substreaked isolate 1 isolates that had not been characterized by WGS. Comparing the *fepR* gene sequence in all 67 substreaked isolates and parent strains revealed that 48/67 of the substreaked *Listeria* isolates in this study showed nonsynonymous mutations in *fepR* (24 missense mutations, 16 nonsense mutations, and 8 frameshift mutations) compared to their respective parent strains (Figure 1). Among the frameshift mutations detected, seven were the result of single nucleotide deletions, and one was the result of a duplication of 10 nucleotides in *fepR* in substreaked isolate FSL H9-0115 (Figure 2). Among the missense mutations detected, 16/24 were localized in the N-terminal DNA binding domain of FepR between amino acid residues 1-43; five nonsense and two frameshift mutations were also detected in this region of FepR. None of the 11 parent strains that carried *bcrABC* or *qacH* acquired the *fepR* mutation in their respective substreaked isolates. ANOVA and post-hoc tests revealed that the presence of *fepR* mutation in substreaked isolates was significantly associated with enhanced substreaked MIC (estimated marginal mean MIC of 5.27 mg/L and 4.21 mg/L for isolates that possess *fepR* mutation and do not possess *fepR* mutation, respectively) (P<0.05), while no significant differences in substreaked MIC values were detected across the three types of mutations detected in *fepR* (i.e., nonsense, missense, frameshift) (P>0.05).

**Figure 2.**
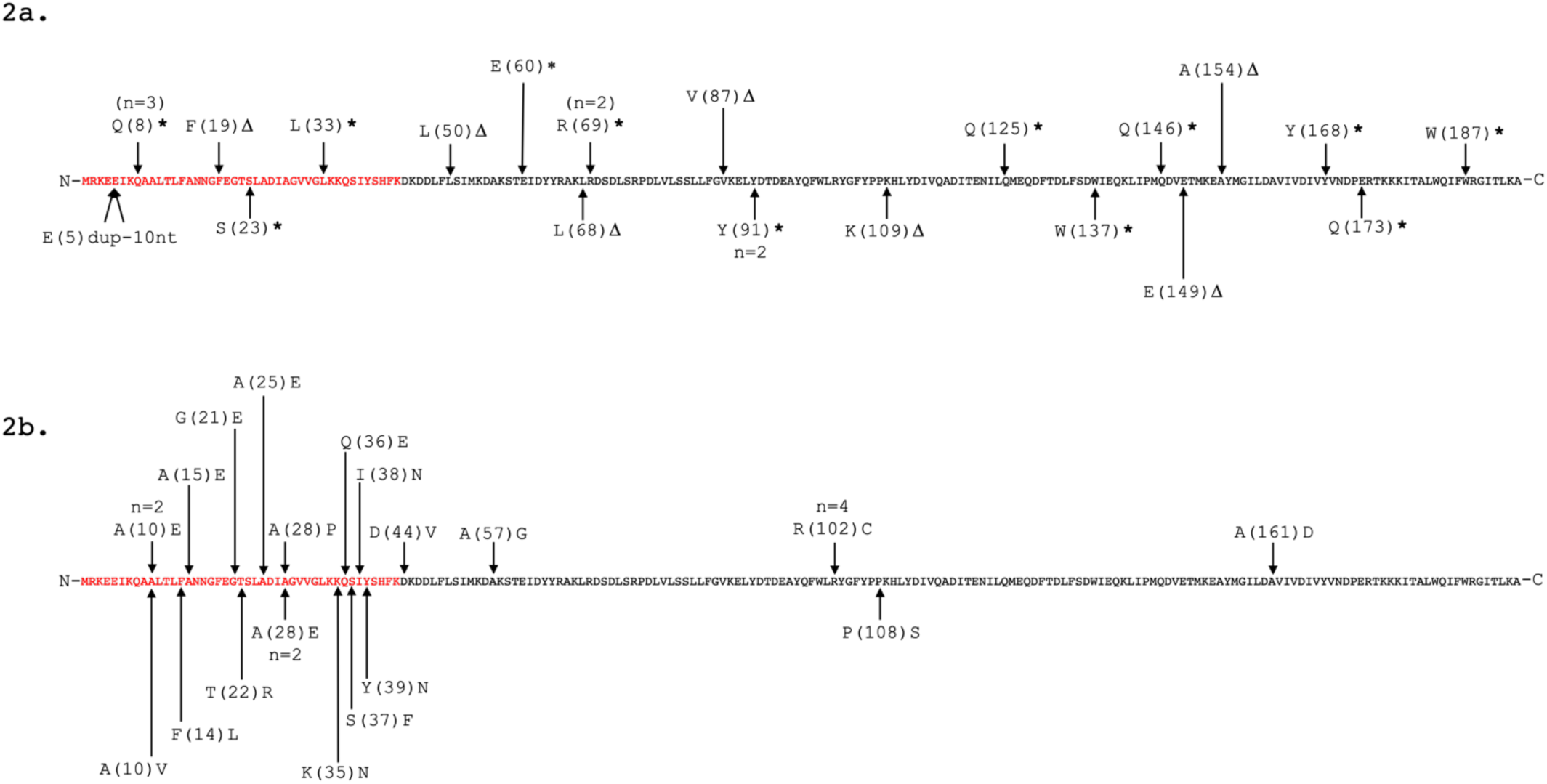
Location of nonsynonymous mutations in FepR in substreaked *Listeria* isolates from this study. Arrows are pointing to amino acid residues that substreaked isolates acquired unique mutations in, and n=number of isolates the unique mutation was detected in. Amino acids colored in red represent the FepR DNA binding domain (43). In figure 2a. nonsynonymous mutations that result in a nonsense mutation are denoted by *, single nucleotide deletions resulting in frameshift mutations are denoted by Δ, and a duplication resulting in a frameshift mutation (denoted by dup-#of nucleotides in length of duplication) (*n*=24) are annotated on the 194 amino acid FepR sequence (GenBank: WP_010991061.1). In figure 2b. nonsynonymous mutations that result in a missense mutation (*n*=24) are annotated on the 194 amino acid FepR sequence (GenBank: WP_010991061.1).

### While some substreaked isolates show loss of *bcrABC*, this does not always result in reduced MIC values

PCR detection of *bcrABC* for substreaked isolates revealed that four substreaked isolates showed loss of *bcrABC* compared to their parent strains (including substreaked isolates 1 and 2 derived from parent strain FSL S11-0256, substreaked isolate 1 derived from parent strain FSL S11-0060, and substreaked isolate 2 derived from parent strain FSL R9-9884). While two of these substreaked isolates showed lower MIC values than their parent strains, the other two substreaked isolates (substreaked isolate 2 from parent strain FSL S11-0256 and substreaked isolate 2 from parent strain FSL R9-9884) both showed the same MIC values as the parent strains (6 mg/L), even though these strains did not show *fepR* mutations, suggesting possible additional mechanisms of adaptation to BC that could be further explored in future studies (Figure 1).

### *Listeria* resistance to low levels of BC is not associated with increased survival in use level concentrations of BC

In addition to investigating *Listeria* resistance to low levels of BC, all parent strains and substreaked isolate 1 isolates were assessed for their survival to a use level concentration of BC (300 mg/L) for 30 s. Initial populations of parent strains (mean of 8.46 ± 0.02 log CFU/mL) and substreaked isolates (mean of 8.45 ± 0.02 log CFU/mL) were reduced by 1.7-6.7 log (mean of 4.56 ± 0.09) and 1.8-6.6 log (mean of 4.48 ± 0.09), respectively, after treatment with BC (Figure 3). Unpaired t-test showed that log reductions of parent strains and substreaked isolates were not significantly different (P=0.57). Furthermore, the 11 parent strains that contained *bcrABC* or *qacH* and the 56 parent strains that did not contain *bcrABC* or *qacH* also did not differ significantly (P=0.65) in observed log reductions in the presence of BC (mean log reductions of 4.47 ± 0.21 and 4.57 ± 0.10, respectively) (raw data deposited in github).

**Figure 3.**
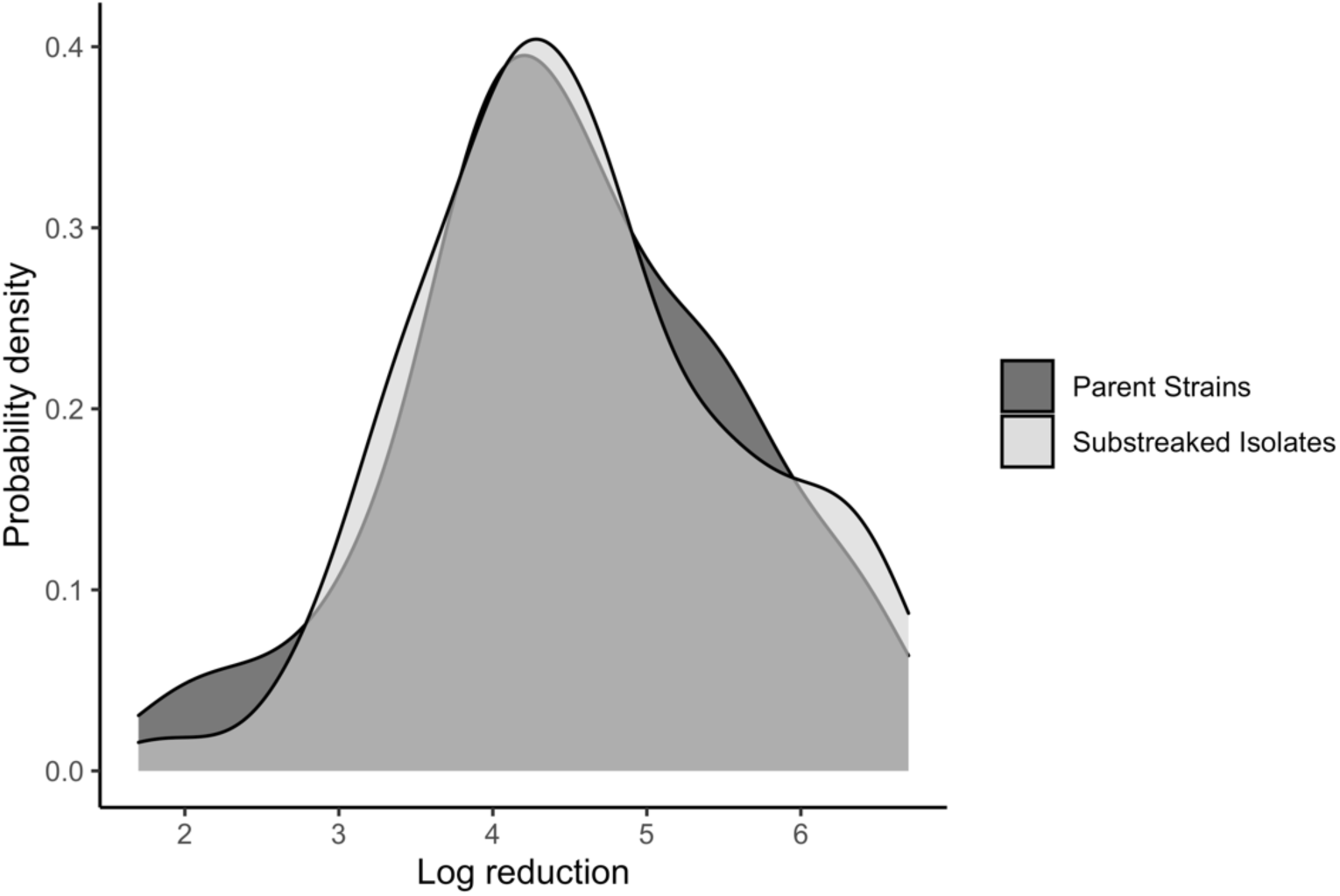
Density plot comparing log reductions of parent strains and substreaked isolates after exposure to use level concentrations of benzalkonium chloride (BC).

### Cocultures of *L. monocytogenes* do not show increased resistance to BC when compared to monocultures

Conjugation experiments with *L. monocytogenes* isolates containing either *bcrABC*, *qacH*, and isolates containing neither genetic resistance determinant, followed by serial passaging in presence of increasing BC concentrations and substreaking of cocultures was performed to determine whether cocultured strains would grow to a higher MIC as compared to monocultures. MIC of cocultures and monocultures immediately after the conjugation experiments on filter plates ranged from 8-10 mg/L, except for the monoculture of strain FSL R12-0334, which showed an MIC of 4 mg/L (Table 6); this was the only monoculture strain that contained neither *bcrABC* nor *qacH*. After serial passaging, all monocultures and cocultures achieved serial passaged MIC in the range of 10-14 mg/L; cocultures did not show significantly higher serial passaged MIC compared to monocultures (P>0.05). Similarly, MIC experiments performed on substreaked isolates obtained from monoculture and coculture serial passage experiments revealed that substreaked MIC across all cocultures and monocultures was 6 mg/L in BC, thus serial passaging cocultures of *L. monocytogenes* in BC did not result in enhanced inherited resistance in substreaked isolates. Aside from determining phenotypic sensitivity to BC, no further characterization of substreaked isolates obtained from coculture serial passage experiments was performed and we did not perform subtyping to determine the specific cocultured strains that were recovered after serial passaging.

**Table 6.** MIC values obtained through three different experiments comparing the MIC of L. monocytogenes monocultures and cocultures to benzalkonium chloride (BC).

| Isolate ID (Shorthand<sup>b</sup>) | lineage, resistance gene | MIC assay<sup>a</sup> |  |  |
| --- | --- | --- | --- | --- |
|  |  | filter plate<br>MIC (mg/L) | serial passaged<br>MIC (mg/L) | substreaked<br>MIC (mg/L) |
| Monoculture<sup>c</sup> FSL S11-0432 (S1) | II, *bcrABC* | 10 | 14 | 6 |
| Monoculture FSL S10-1873 (S2) | II, *bcrABC* | 10 | 14 | 6 |
| Monoculture FSL R9-9884 (S3) | II, *bcrABC* | 10 | 12 | 6 |
| Monoculture FSL R12-0334 (S4) | II, none<sup>e</sup> | 4 | 10 | 6 |
| Monoculture FSL R12-0326 (S5) | I, *bcrABC* | 8 | 14 | 6 |
| Monoculture FSL R12-0189 (S6) | I, *qacH* | 8 | 12 | 6 |
| Monoculture FSL R12-0359 (S7) | I, *bcrABC* | 10 | 14 | 6 |
| Coculture<sup>d</sup> I (S1, S3, S5, S6) | - | 10 | 12 | 6 |
| Coculture II (S1, S5, S6, S7) | - | 10 | 12 | 6 |
| Coculture III (S1, S4, S5, S6) | - | 8 | 10 | 6 |
| Coculture IV (S2, S5, S6, S7) | - | 10 | 12 | 6 |
| Coculture V (S2, S4, S5, S6) | - | 10 | 12 | 6 |
| Coculture VI (S2, S3, S5, S6) | - | 10 | 14 | 6 |

<sup>a</sup>filter plate MIC: Results from MIC assay performed on culture obtained after four *L. monocytogenes* strains were placed onto 0.45 µM filter on BHI agar plates, and allowed to incubate at 22°C for 24 h; serial passaged MIC: the MIC obtained through serial passaging monocultures of cocultures of *L. monocytogenes* in increasing concentrations of BC; substreaked MIC: the MIC value for three isolates that were taken from enumeration plates after serial passage experiments and were substreaked seven rounds onto BHI agar. All three substreaked isolates (substreaked isolate 1, substreaked isolate 2, substreaked isolate 3) displayed the same substreaked MIC.
<sup>b</sup>Isolate shorthand used to describe the four isolates in each coculture.
<sup>c</sup>Individual *L. monocytogenes* strains.
<sup>d</sup>Mixture of four *L. monocytogenes* strains combined and cultured together.
<sup>e</sup>No resistance gene (*bcrABC* or *qacH*) was identified in genome.

## DISCUSSION

In this study, we assessed the capacity of 67 diverse produce-associated *Listeria* isolates to acquire resistance to low levels of BC, a commonly used sanitizer in produce packing and processing environments, and evaluated whether acquired resistance to low levels of BC was associated with increased survival of *Listeria* exposed to use level concentrations of BC (i.e., 300 mg/L). Overall, our results indicate that, despite nearly all *Listeria* isolates showing capacity to acquire inheritable resistance to low levels BC, similar to the level of resistance observed in *Listeria* that carry resistance genes *bcrABC* and *qacH*, this resistance did not significantly impact *Listeria* survival in use level concentrations of BC. These findings provide evidence to support that BC applied to food contact surfaces at typical use level concentrations for quaternary ammonium compounds (e.g., those outlined in US guidance documents, i.e. 21 CFR 178.1010) will have similar effectiveness at controlling *Listeria*, regardless of whether that *Listeria* is resistant to low levels of BC.

### *Listeria* exposed to BC in serial passage experiments acquire inheritable and transient resistance to low levels BC

In this study, we found 105/134 substreaked isolates of *Listeria* were able to acquire and maintain enhanced phenotypic resistance to BC through serial passage in increasing concentrations of BC and seven rounds of substreaking on BC-free media. By showing that this phenotypic resistance could be maintained in substreaked isolates after selective pressure was removed, we have evidence to conclude that the acquired resistance to BC in substreaked isolates was due to inheritance of genetic mutations selected for during serial passaging. This phenomenon is referred to as resistance due to adaption (32). Additionally, we found that all species of *Listeria* represented in this study (*L. monocytogenes*, *L. innocua*, *L. seeligeri*, *L. marthii*, *L. ivanovii*, and *L. welshimeri*) could adapt to levels of BC up to 3-fold higher than their parent MIC. This is consistent with previous studies that have shown 2-3 fold increased MIC to BC in BC-adapted *L. monocytogenes* compared to their wild type MIC (33, 34).

The majority of *Listeria* strains were able to achieve higher serial passaged MIC compared to their parent MIC. However, for many *Listeria* strains this acquired resistance was not maintained or fully maintained after strains were serially-passaged on BC-free media. Thus, serial passage experiments also allowed *Listeria* to acquire transient resistance to BC, which suggests that *Listeria* showed acclimation to BC (32), in addition to adaptation. Several studies have reported Gram positive organisms acquiring transient resistance to BC (35–37), even though some of these studies may not have referred to this observed phenomenon as “acclimation”. For example, in Moore et al. (35) *Staphylococcus haemolyticus* was able to acquire resistance to concentrations of BC 35-fold higher than its parental MIC (MIC value of 0.45 mg/L), but this resistance was lost after passaging in the absence of BC for seven days.

Interestingly, while horizonal gene transfer was previously reported to contribute to dispersal for *bcrABC* (20, 38), in our study cocultures of *L. monocytogenes* strains made up of strains carrying *bcrABC*, *qacH,* or neither genetic resistance determinant did not show evidence for conjugative transfer and selection of *L. monocytogenes* strains with enhanced acquired resistance to BC, as all substreaked isolates obtained from coculture serial passage experiments did not show resistance to concentrations of BC higher than 6 mg/L. While horizontal gene transfer continues to represent a potential mechanism for spread of BC resistance in food processing environments (39, 40), our findings here indicate that horizontal gene transfer may not be a major contributor to the acquisition of BC resistance. However, our study was limited to investigating the horizontal gene transfer among *L. monocytogenes* cocultures, and as such our findings may not be representative of interspecies and intergeneric transfer of BC resistance genes *bcrABC* and *qacH*. For example, interspecies transfer of *bcrABC* has been reported from *Listeria* spp. to *L. monocytogenes* (20), and intergeneric transfer of *bcrABC* has been reported from *L. monocytogenes* to *E. coli* (38).

Notably, our experiments revealed that *Listeria* (cocultures and monocultures) showed adaptation to a maximum MIC of 6 mg/L for substreaked isolates. Similar results were reported in Aase et al. (34), where the authors showed that *L. monocytogenes* isolates with parent MICs to BC ranging from 1-7 mg/L were able to acquire inheritable resistance to a maximum MIC of 7 mg/L BC. Results from Aase et al. (34) and our study both suggest that there is a biological barrier to the level of inheritable resistance *Listeria* can acquire to BC, with that level being ∼6-7 mg/L. Most importantly, these findings support the view that there are limited concerns about *Listeria* acquiring resistance to BC at levels that would impact the efficacy of quaternary ammonium compounds applied at recommended use level concentrations.

### *Listeria* acquires inherited resistance to BC through nonsynonymous mutations in *fepR*

The adaptation mechanism through which the majority of *Listeria* isolates in our study acquired resistance to BC was through nonsynonymous mutations (including missense, nonsense, and frameshift) in *fepR*. Importantly, substreaked isolates representing all *Listeria* species included in this study showed evidence of mutations that would abolish FepR function. FepR has previously been shown to act as a local repressor for transcription of the operon *fepRA* (41). In addition to self-regulation, FepR also represses the transcription of *fepA* (lmo2087), which encodes an efflux pump that is part of the multidrug and toxic compound extrusion (MATE) family (42), which conceivably could remove BC compounds from the bacterial cytosol.

Consistent with our findings, Guérin et al. (41) reported that a single nonsense mutation in *fepR* in *L. monocytogenes* strain BM4716 was associated with a 2-fold higher MIC to BC (MIC value of 8 mg/L), compared to *L. monocytogenes* parent strain BM4715 (MIC value of 4 mg/L). Moreover, strain BM4716 and a *fepR* deletion mutant of BM4715 (BM4715*ΔfepR*) both showed a 64-fold increase in gene expression of FepA as compared to BM4715, demonstrating that loss of function of FepR results in overexpression of FepA. However, MIC assays conducted in the presence of the efflux pump inhibitor reserpine did not lead to the expected reduction in MIC in strain BM4716 (41), which could be interpreted as suggesting that overexpression of efflux pump FepA may not be responsible for the enhanced BC resistance associated with *fepR* mutations. Alternatively, reserpine added at a concentration of 10 mg/L may not have been used at the appropriate concentration to inhibit a highly expressed FepA or may not inhibit FepA as effectively as it does other efflux pumps; this is consistent with observations in Meier et al. (18) where the authors found that 10 mg/L reserpine did not inhibit efflux activity in all eight *Listeria* strains that carried *bcrABC* in their study (18). Overall, data available to date strongly support that mutations in *fepR* resulting in truncation and loss of function in FepR are associated with and responsible for resistance to low levels of BC.

In addition to mutations resulting in truncation of FepR (nonsense and frameshift mutations), we also observed missense mutations in 24 substreaked *Listeria* isolates, and 22 of those isolates also showed the phenotype of acquired enhanced resistance to BC. Similar to our findings, Bland (43) showed that *L. monocytogenes* can acquire similar levels of resistance to BC through both nonsense and missense mutations in *fepR*. In our study, we detected the majority (16/24) of missense mutations in the N-terminal DNA binding domain (NDB, helices *α*1 to *α*3) of FepR, which makes up the helix-turn-helix (HTH) motif that binds to the operator sequence of DNA on the *fepRA* operon (44). This NDB region of FepR is highly conserved with other TetR family transcriptional regulators (44), hence previous findings in FepR homologues may be used to elucidate the likely effect of these missense mutations on FepR function. For example, TetR family regulators QacR and TetR both show a lack of water present at the NDB-operator DNA sequence binding interface, which facilitates tight binding of their NDB regions to DNA operator sequences (45, 46). Here, we saw 8/16 mutations in the NDB region of FepR that showed an amino acid change to glutamic acid (E) and 3/16 mutations that showed an amino acid change to asparagine (N), both of which have polar side chains that can participate in hydrogen bonding. We hypothesize that the incorporation of E and N residues in FepR’s NDB allows for the incorporation of water into the NDB-operator DNA sequence binding interface of *fepRA*, which can cause reduced or inhibited binding of FepR to the *fepRA* operator.

In addition, in our study we also observed a handful of missense mutations located in regions localized outside of FepR’s NDB. One such mutation that was observed in *L. innocua* FSL H9-0107, a substreaked isolate that showed 2-fold enhanced resistance to BC compared to its parent strain, resulted in a change of amino acid from Proline to Serine at residue 108. Notably, this same amino acid change was reported in Bland (43) in *L. monocytogenes* strain WRLP380, a strain which conferred 1.5-fold enhanced resistance to BC compared to its parent strain. Together, the findings from Bland (43) and our study demonstrate that this particular amino acid change is associated with FepR loss of function, and further investigation is warranted to uncover the mechanism responsible for this observed phenotype. Additionally these findings highlight the ability of *L. monocytogenes* and *L. innocua* to acquire BC resistance through the same nonsynonymous mutation in *fepR*, further emphasizing the validity of *L. innocua* as an index organism for the presence of *L. monocytogenes* in food processing environments where BC is used for sanitation (47). Interestingly, however, no *fepR* mutations were present in parent strains of *Listeria* in this study, suggesting selection against the loss of function mutation in *fepR* in more complex environments. Hence, the practical implications of *fepR* mutations in natural environments remains to be determined.

### BC resistance phenotype is similar across *Listeria* with efflux mechanisms mediated by *bcrABC* and *qacH* or *fepR* mutations

The presence of *bcrABC* and *qacH* in *Listeria* strains from this study was associated with higher phenotypic resistance to low levels of BC (parent MIC values of 4-6 mg/L) compared to *Listeria* strains that did not carry these genes (parent MIC values of 1-2 mg/L). Similar findings have been reported for isolates carrying *bcrABC* and *qacH* obtained from a variety of food processing environments (9, 16, 18). For example, one study (9) reported that *Listeria* isolates carrying *qacH* showed MIC values ranging from 5-12 mg/L (as compared to strains without *qacH* that showed MIC values of ≤ 5 mg/L), and Cooper et al. (16) showed that MIC values for *Listeria* isolates carrying *bcrABC* were 2.5-3.5 fold higher than *Listeria* isolates that did not possess a BC resistance gene.

Notably, none of the substreaked isolates in our study that were derived from parent strains carrying *bcrABC* or *qacH* acquired a mutation in *fepR*. Moreover, no *Listeria* with any of these three genotypes associated with BC resistance (presence of *bcrABC* or *qacH*, or nonsynonymous mutation in *fepR*) showed substreaked MIC values >6 mg/L. Because there does not seem to be any phenotypic advantage in having one of these three genotypes over the others, we hypothesize there was a lack of selective pressure for *Listeria* strains already carrying either *bcrABC* or *qacH* to acquire inheritable resistance through nonsynonymous mutations in *fepR*.

### *Listeria* resistance to low levels of BC does not correlate with increased survival in use levels of BC

While the majority of *Listeria* isolates in this study were able to acquire, through serial passage experiments, resistance to low levels of BC at a biological barrier of 6 mg/L, when we compared the survival of isolates with acquired resistance to low levels of BC to their respective parent strains at a use level concentration of 300 mg/L BC, the substreaked isolates with acquired resistance to low levels of BC did not show better survival compared to parent strains. These findings are in accordance with Kastbjerg and Gram (48) who also found that *L. monocytogenes* that were adapted to grow in 48 mg/L of BC did not show better survival in 125 mg/L BC compared to unadapted *L. monocytogenes* parent strains.

Additionally, the 11 parent strains carrying *bcrABC* and *qacH* also did not show increased survival after exposure to 300 mg/L BC compared to the other 56 parent strains in this study. In Cooper et al. (16), the authors suggest that the presence of BC resistance genes, specifically *bcrABC*, can represent indicators for *Listeria* persistence in food processing environments. Based on our results, the presence of *bcrABC* confers *Listeria* resistance to BC, but at levels well below those applied at use level, and *bcrABC* does not confer increased survival of *Listeria* upon exposure to use level concentrations of BC. Therefore, based on our results it is unlikely that presence of *bcrABC* would represent a plausible indicator of *Listeria* persistence in food processing environments.

## Acknowledgement

We want to thank Dr. Yi Chen at FDA-CFSAN for kindly providing isolates FSL R12-0098 (SRR1027706), FSL R12-0099 (SRR1101447), FSL R12-0085, and FSL R12-0093. We also want to thank Dr. Haley Oliver for kindly providing isolates FSL R12-0180 (SRR9019233), FSL R12-0260 (SRR9019214), FSL R12-0324 (SRR9019309), FSL R12-0133, FSL R12-0181, FSL R12-0189, FSL R12-0326, FSL R12-0334, and FSL R12-0359. The work upon which this project entitled “*Listeria* develops reduced sanitizer sensitivity but not resistance at recommended sanitizer use levels” was funded, in whole or in part through a subrecipient grant awarded to The Center for Produce Safety through the Florida Department of Agriculture and Consumer Services Specialty Crop Block Grant Program. Any opinions, findings, conclusions, or recommendations expressed in this publication are those of the author(s) and do not necessarily reflect the view of The Center for Produce Safety and the Florida Department of Agriculture and Consumer Services.

